# Gist – an ensemble approach to the taxonomic classification of metatranscriptomic sequence data

**DOI:** 10.1101/081026

**Authors:** Samantha Halliday, John Parkinson

## Abstract

The study of whole microbial communities through RNA-seq, or metatranscriptomics, offers a unique view of the relative levels of activity for different genes across a large number of species simultaneously. To make sense of these sequencing data, it is necessary to be able to assign both taxonomic and functional identities to each sequenced read. High-quality identifications are important not only for community profiling, but to also ensure that functional assignments of sequence reads are correctly attributed to their source taxa. Such assignments allow biochemical pathways to be appropriately allocated to discrete species, enabling the capture of cross-species interactions. Typically read annotation is performed by a single alignment-based search tool such as BLAST. However, due to the vast extent of bacterial diversity, these approaches tend to be highly error prone, particularly for taxonomic assignments. Here we introduce a novel program for generating taxonomic assignments, called Gist, which integrates information from a number of machine learning methods and the Burrows-Wheeler Aligner. Uniquely Gist establishes the most appropriate weightings of methods for individual genomes, facilitating high classification accuracy on next-generation sequencing reads. We validate our approach using a synthetic metatranscriptome generator based on Flux Simulator, termed Genepuddle. Further, unlike previous taxonomic classifiers, we demonstrate the capacity of composition-based techniques to accurately inform on taxonomic origin without resorting to longer scanning windows that mimic alignment-based methods. Gist is made freely available under the terms of the GNU General Public License at compsysbio.org/gist.

## INTRODUCTION

Recent advances in high-throughput sequencing are driving new programs of research that are profoundly transforming our understanding of the relationship between microbiomes and their environments, including enteric bacterial communities with significance for human health. Typically, studies which rely on 16S rRNA surveys and metagenomic DNA analyses yield only limited insights into gene activity. To address this, whole-microbiome gene expression profiling (‘meta-transcriptomics’) through RNA sequencing, has emerged as a means of gaining a mechanistic understanding of the complex functional relationships within microbial communities.^1^ Metatranscriptomic data have been collected for ecosystems as diverse as deep sea hydrothermal vents^2^, forest soils,^3^ and both animal^4,1^ and human^5^ body environments.

Due to the size of the datasets and relatively short sequence lengths, the processing, annotation, and interpretation of metatranscriptomic data are all significant technical challenges. Of particular interest is the ability to accurately assign taxonomic information to individual sequences, which yields insights into the contribution of individual taxa to biological processes within the microbiome. For example, obesity-associated inflammation (shown to cause insulin resistance) is thought to be driven by induction of a low-grade inflammatory response in the intestinal epithelium by Clostridial species normally suppressed by the TLR5 immune pathway;^6^ however a detailed mechanistic understanding is lacking. Similarly, key fermentation products of abnormal bacterial metabolism in the human gut (short-chain fatty acids, especially propionic acid) produced by certain *Clostridia, Desulfovibrio*, and *Bacteroidetes* species, have been shown to trigger neuroinflammation in a mouse model, resulting in behavioral changes consistent with autism spectrum disorders.^7^ Further mechanistic insights into the molecular basis for disease pathogenesis may be gained through the application of metatranscriptomics, identifying differentially expressed genes and pathways associated with the disease state. Additionally, precise taxonomic annotations permit these pathways to be attributed to discrete taxa.

Beyond understanding the biochemical contributions of individual taxa within a complex community, the taxonomic classification of raw sequence reads has the potential to streamline sequence assembly for both metatranscriptomes and metagenomes.^8^ *De novo* assembly (assembly without a scaffold) is typically performed using techniques that are computationally intensive and scale poorly with respect to read count.^9, 10^ By binning reads according to species labels, the number of necessary comparisons can be significantly reduced, simultaneously minimizing the generation of chimeric contigs.

Previous studies concerned with taxonomic profiling of microbiomes have largely focused on analysis of well-characterized 16S ribosomal RNA fragments, often using unusual combinations of methods to achieve high accuracy.^11, 12, 13, 14, 15^ Here, our aim is to develop a framework that extends this classification quality to all genes. Three general categories of techniques for assigning taxonomic labels have been developed: phylogenetic, alignment-based (sometimes called ‘similarity-based’), and compositional.^15^ Phylogenetic strategies, which emphasize a structural understanding of the underlying evolutionary tree,^15^ can be thought of as an extension to either of the other two, and involve the reprocessing of results from alignment and/or composition based methods to quantify the distance between the assigned reads and the reference data. In alignment-based strategies, the results of a search method such as BLAST^16^ are used to map reads directly onto known reference sequences (e.g. genomes). Due to the reliance on databases that represent only a fraction of bacterial diversity, these methods perform poorly for data containing taxa that have not previously been well-sampled.^17^ Horizontal transfer events are especially likely to cause erroneous assignments.^11^ When dealing with unknown sequences, compositional approaches are more robust. In most implementations these rely on counting the frequencies of short fragments of reads, called *k*-mers, using a sliding window of some preset length *n* to scan either the nucleotide or amino acid content of a given reference sequence. These counts are then tallied into a histogram which provides a position-independent summary of the sequence content. Reads as short as 35 nt have been seen to exhibit codon bias,^11^ relative nucleotide abundances, and other features that can be recognized easily with such histograms. Popular machine learning techniques for this problem include naïve Bayes (NB),^11^ *k*-means,^13^ hidden Markov models (HMM),^18^ and Gaussian-kernelized *k*-nearest neighbors (*k*NN).^12^

Here we present a new ensemble classifier, Gist, developed specifically to exploit the information richness of metatranscriptomic data. Uniquely Gist establishes weights for each technique that are specific to each genome in its reference dataset, allowing for the achievement of high-quality results using much smaller *k*-mers than in previous implementations of composition-based approaches. We validate our approach using a novel synthetic data simulation pipeline, Genepuddle, as well as data from a previous study of the intestinal microbiome associated with a non-obese diabetic (NOD) mouse model. The performance of Gist is compared against three other metagenomic compositional classifiers (KRAKEN,^19^ CLARK,^20^ and NBC^11^). With its improved performance and reduced reliance on comprehensive genomics databases, we propose Gist as an effective solution to the taxonomic classification of metagenomic and meta-transcriptomic datasets.

## RESULTS

### Gist – an ensemble taxonomic classifier for analyzing metatranscriptomic sequence datasets

Here we present Gist (Generative Inference of Sequence Taxonomy), an ensemble classifier that combines the output of the Burrows-Wheeler Aligner (BWA)^21^ and four statistical models used for examining *k*-mer composition: naïve Bayes (NB), 1-nearest neighbor (1NN), a Gaussian mixture model (GMM), and a novel technique, the expected co-delta correlation (ECC) to assign taxonomic labels to metatranscriptomic read data (Figure 1).

BWA was chosen primarily to provide the program with an efficient method for finding close or exact matches. Like many mapping aligners, BWA has a low tolerance for noise, which significantly curtails the rate of false positives. BWA was selected over other programs of its type due to its low memory usage and high accuracy when analyzing prokaryotic sequences.^22^ We avoided the use of protein alignments in Gist due to their propensity of locating paralogs, limiting accurate taxonomic assignment.

Naïve Bayes, the most popular algorithm for compositional taxonomic classifiers, works by assuming the data is distributed according to a single multivariate Gaussian distribution. It is effective at determining the mode of the distribution (i.e. what typical genes from a genome look like), but does not perform well on outliers, and rapidly becomes oversaturated with smaller *k-*mer sizes. It may also fail when a genome has several large, distinctive subpopulations, causing the predicted mean to fall into an area of low importance.

*K*-nearest neighbors is an instance-based method that determines the nearest known *K* genes, like an aligner, in each species and reports the distance. It provides some flexibility over the BWA component as it is more tolerant of rearrangements and short duplications, although it is sensitive to missense mutations and can be confounded by certain anagrams of repetitive sequences. *K* = 1 was chosen to minimize interference from large families of paralogs.

A Gaussian mixture model (GMM) is a derivative of NB which uses expectation–maximization to find the means and variances of multiple subpopulations of genes. It is most useful when the genome is not well-modeled by one distribution, such as when it has recently undergone large-scale horizontal gene transfer from another source, has many genes that do not obey normal codon distributions (e.g. RNA genes), or if it contains a large family of proteins with many paralogs. Expectation–maximization is a randomized iterative algorithm, however, and therefore much iteration is required on average to attain a model of acceptable quality.

Expected co-delta correlation (ECC) is a novel technique that provides an efficient tradeoff to the calculation of full covariance tables for Gaussian-based methods. It calculates the rates of co-occurrence between pairs of *k*-mers within the read, and then compares this to the average rates for each genome. Because of this two-dimensional relation, ECC can encode motifs of longer lengths by connecting *k*-mers found together in one gene to each other, even if these are discontiguous; e.g. when used with translated protein sequences, it can determine the amino acids most commonly found adjacent to disulfide bridges.

While each component excels at identifying key features representative of specific genomes, individually they are too simplistic to model the *k*-mer landscapes necessary for accurate classification. For example, ECC does not consider the background rate of each *k*-mer and must therefore be combined with other techniques to be effective. Consequently, a single-layered neural network, analogous to logistic regression, is employed to determine the best combination of methods to describe each genome. This approach significant improves resolution power at short *k*-mer lengths compared to existing composition-based methods^17^ operating under the same constraint; the resultant joint distribution more accurately represents the shape of each genome’s total gene population, and can better predict the expected compositional signatures of unknown genes from related strains, in large part due to the short *k*-mer lengths employed. More details of the implementation of each method can be found in Supplemental Methods. Generating the expected weights for each technique is performed during an initial bootstrapping process that is tailored to the dataset provided to the program. This requires synthetic data drawn from a distribution approximating the real reads. To generate such data, we created a novel prokaryotic meta-transcriptome simulation pipeline, Genepuddle. Obtaining the underlying distribution of taxa can be accomplished using 16s rRNA counts, and the algorithm is robust enough to permit the substitution of genomes from other genera and families when more precise species data is not available. (See Supplemental Methods.) While constructing a generic weight and class set is feasible, narrowing the list of possible assignments greatly reduces sequence space noise and improves overall accuracy.

Integration of classifier results to yield overall probabilities that a read derives from a given genome is accomplished with Bayesian inference: the log-scores from each component are weighted with the information obtained from training the neural network, and then summed. For a single read, this process is repeated for every genome in the dataset. The final output report is generated using a two-pass method which ensures that the program returns larger taxonomic units (i.e., less precise predictions) if the best-scoring taxon appears to be drawn from the same pool as its immediate relatives when subjected to a one-tailed *t*-test. This ensures that unique mutations are correctly assigned to their source strain, while highly diverse genes are assigned to the parent species or even genus.

### Comparison of Gist against other taxonomic classifiers using a simulated dataset

To assess the reliability of any classifier, it is valuable to have a source of data for which the correct labels are known. Often when developing metagenomic methods, the MetaSim package^23^ is used to generate simulated reads for this task, but this does not consider the types of fidelity loss produced during RNA sequencing and so is inappropriate for simulating a meta-transcriptome. We created a pipeline for this, **Genepuddle,** which prepares genomes from the NCBI site for use with Griebel *et al.’s* Flux Simulator,^24^ allowing rapid and efficient production of large sets of metatranscriptomic reads according to a specified error model, read length, and abundance distribution.

For this study, a set of 295 genomes was examined, belonging to genera and families associated with the NOD mouse dataset described further in the methods section. Two synthetic datasets were created for testing: an *unbiased* version, with equal counts for each strain, and a *biased* version, with abundances derived from 16S data with some adjustment, to better reproduce the relative levels expected in real data. (See Supplementary Notes.)

The results were scored using another new tool called **Lincomp,** which reports accuracy at the smallest matching taxonomic rank; e.g. if a classifier guesses the wrong species in the correct genus, then it is considered to be reliable at the genus level for that hit. Lincomp uses the NCBI taxonomy.

Training consisted of 10,000 reads per strain, totaling 2,950,000 reads. Unbiased and biased test datasets consisted of 737,500 reads (2,500 per strain) and 85,990 reads, respectively. **Figure 2** shows the precision of each program on these datasets with variable levels of isotropic noise (randomly-replaced nucleotides) to simulate biological diversity despite using a limited set of curated reference genomes. As reference genomes are rarely representative of sequences found in the wild, it is necessary to emphasize experiments which do not disproportionately reward programs that focus on exact matching.

On data with no noise, NBC’s use of 15-mers presents an effective method for matching the presented synthetic data with the corresponding sequences in the database. KRAKEN and CLARK both aim to improve on NBC’s running time by database pruning and through the clever use of hashing methods. While CLARK maintains excellent precision, both of these approaches decline in performance proportional to the level of noise present in the data, as their comparatively long and fragile *k*-mers (both default to 31 nt) are interrupted by mutations. Gist’s performance curves are closer to those of NBC, showing strong resilience to noise, as predicted by its use of much smaller *k*-mers in its ensemble.

### Gist’s algorithmic approach combines several different methods using per-genome weights

Figure 3 illustrates some of the model parameters learned by Gist during its training process.

Part (A) shows the weights learned by the Autocross training algorithm for the simulated mouse data in Figure 2. During classification, each of the nine methods shown generates a score, which is multiplied by the corresponding weight and summed to produce the final class-read score. These weights thus reflect the contrastive balance necessary to distinguish each genome’s reads, as well as a representation of how well each element of the ensemble models each genome. The weights are shown normalized, as the genomes have highly variable total weight (from 10^−5^ to 10^−1^.)

Part (B) illustrates part of how the profiles of genomes are represented by the peptide naïve Bayes classifier. In this example, the average frequencies of different amino acids in six genomes *(Lactobacillus crispatus* ST1, *Streptococcus agalactiae* A909, *Clostridium difficile* 630, *Microluntaus phosphovorus* NM-T^T^, and *Escherichia coli* K-12, substr. MG1655) are compared, highlighting that, for example, the peptide dimer SerAla is less common in the Firmicutes shown than in *E. coli* or *M. phosphovorus*, but LeuLys is more common. *L. crispatus* ST1 and *S. agalactiae* A909 are both from the same family of Firmicutes (Lactobacillales) and are hence both related to *C. difficile* 630, another Firmicute, but more closely to each other. *M. phosphovorus* NM-1^T^ is included as an out-group, and is from the phylum Actinobacteria.

Part (C) shows expected codelta tables for the peptide dimers in *S. agalactiae* A909, *L. crispatus* ST1, and *C. difficile* 630. Visually, these strongly resemble one another, featuring several strong disequilibrium bands, although not all are common to all three graphs. Additionally, it can be seen that Firmicutes emphasize Isoleucine and rely less on Glutamine and Asparagine than any of the other strains shown in part (D).

Part (D) shows the contrast between genomes and random noise generated with the same read length distributions and GC content. The closer the genome is to random noise, the less likely the identities of its individual amino acids matter, creating a simple test metric for genome functionality by comparing similarities. *Carsonella ruddii* and *M. phosphovorus* are two very atypical genomes; *C. ruddii* is an endosymbiont found in psyllids (jumping plant lice) with a genome of only 160 kilobases^25^ and *M. phosphovorus* is a chemoorganotroph notable for lacking several pathways typical of other Actinobacteria.^26^

### Comparison of Gist against other taxonomic classifiers using real metatranscriptomic datasets

To ascertain the performance of Gist on real data, a model dataset was used: the mouse colon data mentioned previously. The data used in this experiment consisted of three samples, identified as 501, 502, and 504, which were collected from the mice’s colons and chemically treated to delete rRNA as described in Xiong, Frank, *et al*. 2012, then sequenced using the Illumina platform^1^. The mice were reared under germ-free conditions and initially colonized with altered Schaedler flora (ASF), a community of 9 strains of bacteria commonly found in the murine gut as described in Table 2. The objective was to demonstrate how 16S data can be used to bootstrap the evaluation process, so that it is not necessary to compare metatranscriptomic data against all known taxa, which exaggerates the risk of misclassification.

**Table 1.** Taxa in the simulated mouse dataset.

| Genus | Strains |
| --- | --- |
| Actinomadura | 3 |
| Aerococcus | 2 |
| Anaerostipes | 3 |
| Bacteroides | 9 |
| Basfia | 1 |
| Blautia | 2 |
| Brevibacillus | 6 |
| Brevibacterium | 2 |
| Butyrivibrio | 2 |
| Dorea | 3 |
| Enterococcus | 11 |
| Eubacterium | 7 |
| Exiguobacterium | 3 |
| Glaciibacter | 1 |
| Idiomarina | 5 |
| Lachnoanaerobaculum | 2 |
| Lactobacillus | 52 |
| Leifsonia | 2 |
| Mannheimia | 3 |
| Microlunatus | 1 |
| Moorella | 1 |
| Mucispirillum | 1 |
| Paenibacillus | 9 |
| Parabacteroides | 1 |
| Pelotomaculum | 1 |
| Peptostreptococcus | 3 |
| Propionibacterium | 14 |
| Roseburia | 5 |
| Staphylococcus | 53 |
| Streptococcus | 79 |
| Thermaerobacter | 2 |
| Thermomonospora | 1 |

**Table 2.** Taxa in the Altered Schaedler Flora (ASF).

| Taxon ID | Name |
| --- | --- |
| 97138 | **Clostridium** *sp.* ASF356 |
| 97137 | **Lactobacillus** *sp.* ASF360 |
| 1235801 | **Lactobacillus murinus** ASF361 |
| 1379858 | **Mucispirillum schaedleri** ASF457 |
| 1235802 | **Eubacterium plexicaudatum** ASF492 |
| 1378168 | **Firmicutes bacterium** ASF500 |
| 84086 | *unclassified Firmicutes sensu stricto (miscellaneous)* |
| 97139 | **Clostridium** *sp.* ASF502 |
| 1235803 | **Parabacteroides** *sp.* ASF519 |

16S processing of this dataset produced a large inventory of candidate matches of diverse species within the orders of the expected strains; samples from the top 25 genera, plus one clade known to be present but not represented in the 16S data, were included in the database for this problem. The result was a dataset consisting of a total of 295 strains taken from the NCBI FTP server, many of which were still in the draft or assembled contig stage of processing at the time of collection. Training and test data were generated using Genepuddle, and abundances were then calculated for the reads (from the 501, 502, and 504 datasets). The results of classification are summarized in part B of Figure 4.

In order to evaluate the performance of these models on the real data, the combined outputs of three aligners (BWA, BLAT, and BLAST) were used with the true ASF genomes (Table 2) to produce a ‘gold standard’. The Lincomp tool was then used to evaluate the accuracy of each method with respect to this gold standard. The results are shown in Figure 4, part A.

In the case of the NOD mouse dataset, the rRNA removal treatment appears to have been biased against the phylum *Bacteroidetes*, resulting in the complete removal of *Parabacteroides* from the data. This resulted in a deficiency in the constructed database which was amended by adding in the taxa by hand. This illustrates the importance of careful 16S curation, and how rRNA removal products can be biased towards certain taxa.

CLARK and KRAKEN obtained similar levels of performance at the strain level to Gist and NBC, but were only able to classify a very small portion of the data (5-10%) in each test case. Conversely, NBC and Gist exhibited much greater success in identifying related sequences among the 293 strains present in the database which were not identical to those in the ASF data.

## DISCUSSION

Here we have presented a novel taxonomic classification algorithm, Gist, which yields precision and sensitivity on-par or surpassing existing composition-based methods at a very short *k*-mer length by combining the outputs of a number of simple techniques. This enables Gist to overcome the robustness problems that characterize programs such as NBC and have led to widespread disuse of *k*-mer based methods for classifying gene sequences taxonomically.

Bazinet and Cummings^15^ found in 2012 that the Naïve Bayes Classifier (NBC)^11^ yielded the best performance amongst compositional techniques.^15^ NBC uses only a single nucleotide Naïve Bayes distribution to classify reads. To function with such accuracy comparatively large *k-*mers are required, but the recommended default, 15 nt, can be defeated entirely with a pathological example containing only 6.7% noise, where every fifteenth nucleotide has been corrupted. This is an issue frequently mentioned^27–29^ in the literature surrounding spaced seeds, also known as gapped *k*-mers, which are usually discussed in the context of alignment methods. Both KRAKEN and CLARK have spaced seed implementations, SEED-KRAKEN^30^ and CLARK-S.^31^ These have yielded mixed results, with KRAKEN showing substantial improvements, but CLARK showing little increase in sensitivity, perhaps due to conflicts with its database pruning methodology, which eliminates much of the redundancy that might otherwise serve to make the program resilient to small mutations.

Most compositional classifiers for metagenomic taxonomy implement only one machine learning technique: RITA^17^, NBC^11^, KRAKEN,^19^ and CLARK^20^ are all examples of NB-based programs, TACOA^12^ uses *k*NN, MetaCV^32^ uses a modified protein-based HMM, and Phymm/PhymmBL^18^ uses a collection of HMMs called an interpolated Markov model. The most popular short-fragment taxonomic classifier to use more than one compositional technique is RDP^33^, which uses NB and *k*NN, but it is intended for use with ribosomal RNA fragments. (NBC, TACOA, and PhymmBL are reviewed further in Bazinet and Cummings^15^ along with many other methods for taxonomic classification, their relative performances, and strategies.) The multifaceted approach used by Gist makes it possible to combine the probability spaces generated by each classifier into a unified model where different elements can be emphasized in order to better fit the peculiarities of the individual genomes, as illustrated in Figure 3, part A. This gives Gist much more flexibility in accommodating the complexity of these distributions without succumbing to the type of over-fitting one would expect from a strictly instance-based technique such as pure *k*NN while avoiding the cost of training a powerful discriminatory method such as a support vector machine to accommodate every new classifier category.

Key to the difficulty in using simpler classifiers with this type of data are the many unique limitations held by each method. For example, even when only considering a well-behaved distribution of nucleotides with a single mode, the "naïvety" of Naïve Bayes poses a conceptual problem for compositional classification. The coordinates of each data point (i.e. the histogram of each read) must sum to 1, because they are defined fractionally, and the use of a sliding window means that the value of each successive *k*-mer is highly dependent on the value of the previous *k*-mer. Ignoring such correlation is an attractive optimization for such large datasets, but it hinders discrimination between closely-related strains. In Gist’s implementation of 1NN, a correction is applied to accommodate for this same problem; each read and gene is normalized by its length during scoring, resulting in distances that do not sum to 1, but reducing the gap between the correct gene and its reads. To deal with this defect in Gaussian methods, expected codelta correlation is added, allowing correlation tests to be performed on unordered high-dimensional data (unlike e.g. Pearson correlation) with samples containing only one member (unlike direct covariance testing.)

Previously, generic taxonomic classifiers also failed to consider the wild variability of branch length on the tree of life: three taxa of equivalent rank with the same parent are unlikely to have the same sequence distance from each other. While classifiers including RITA have looked at setting constant thresholds for taxonomic units according to the user’s intuition,^17^ the developers ultimately assumed that each taxon had a flat evolutionary distance from its siblings defined only by its rank, a generalization that is poorly supported by even the most cursory survey; the authors are aware of no program that attempts to learn class boundaries for short reads for the purpose of intuiting taxonomic thresholds. Gist elides the need for this as a consequence of the Autocross training process, where contrasts between different taxa are maximized to ensure as many sequences as possible in the training set are classified correctly, as illustrated in Figure 3a. The decision of whether to assign a hit to a parent or child taxon is made solely on the consensus of the children (i.e. the read could be assigned equally to any of them), not on any assumptions about branch length. Given the inconsistent and sometimes discontiguous^34^ nature of bacterial evolution, it would appear to be critical that, in lieu of a more orthogonal alternative to Linnaean taxonomy, methods prioritize ways to work around it.

As noted by the authors of NBC,^11^ 16S data will not always be attainable for a dataset— or discernible for closely-related environments. During the development of Figure 2, we found that Gist was equally successful at classification regardless of whether or not it was trained with data of uneven abundance. It is expected that this result will scale to a universal weight set, i.e. one which includes all available sequenced strains, although the compositional elements of the method may face challenges due to the overcrowding of small *k*-mer space, an obstacle also noted by NBC’s authors.

This result also reveals something about the prominence of compositional features—in the biased training dataset, the weights of some strains were derived from as few as 4 reads of 100 nt in length each. This amount of sequence data is nowhere near sufficient to provide complete coverage for most bacterial coding sequences, and yet training was still highly successful and did not lose significant sensitivity on the test dataset. Genus-level pooling is partly to be credited for this, e.g. *Streptococcus* was backed by 80 strains, totaling 423 sequences.

From a statistician’s perspective, Gist is a relatively simple ensemble; while perhaps more sophisticated than existing taxonomic classifiers, it is nowhere near the complexity of the monstrous ensemble models that cracked the 2009 Netflix prize. Despite this, however, Gist is able to achieve considerable performance as a result of the architectural choices that it showcases. These features—the use of supervised learning on simulated data, the combination of different compositional classification techniques, and the use of an unpaired *t*-test in phylogenetic comparison—can readily be applied to different methods (and even different problems.) The possibilities of building assumptions about relative abundance into a metagenomic database also show promise (and pitfalls) as a way to eliminate artifacts of the classification process.

Finally, Gist does not yet support the use of RPKM (reads per kilobase per million) or any other measure to normalize the weights of genes during initial class construction; they are regarded as having an equal chance of being hit against, regardless of length. Adding RPKM will improve Gist’s performance further.

Taxonomic classifiers, like many kinds of bioinformatics programs, have a relatively poor survival rate in the wild. Many (such as MetaCV, NBC, and RITA) cease development shortly after publication. One unfortunate consequence of this is that the databases offered with these programs become outdated, as in the case of NBC’s webserver (not updated since March 2011) and MetaCV’s cvk6 database, which has not been updated since October 2012 despite its website’s assurances of continuing monthly updates and a January 2013 release of the program itself. Given this pattern, it is tempting to speculate that many estimates of species diversity based on gene sequencing could be grossly exaggerated, as *k*-mer and alignment methods lacking a minimum threshold criterion for classification may report random, unrelated taxa based on very small amounts of evidence.

To ensure that Gist does not become obsolete due to changes in reference databases, we are currently developing an efficient pipeline for updating and constructing both databases and training data tuned to the latest available information and the user’s needs. In addition, with Gist’s Bayesian inference framework for coordinating the outputs of its classifiers, it should be practical to integrate other open-source sequence classification programs also, ensuring a reliable source of high-quality read classifications for years to come.

## ACKNOWLEDGEMENTS

This work was funded by grants from Genome Canada and Natural Sciences and Engineering Research Council of Canada (RGPIN-2014-06664). High performance computing was provided by the University of Toronto SciNet facility. We would to thank Dr. Richard Zemel (University of Toronto Department Computer Science), Dr. Anna Goldenberg (Genetics and Genome Biology, Hospital for Sick Children) and Dr. Michael Brudno (University of Toronto Department of Computer Science, *et alia)* for valuable discussions in developing the Gist algorithm.

## FIGURE LEGENDS

**Figure 1. Program overview.** A) In a simplified example of the *k*-mer construction process, the tallies of each 3-mer are treated as the coordinates of a 3^4^-dimensional vector. B) Usage of the Gist and Genepuddle pipelines to analyze a metatranscriptome for which 16S data is available.

**Figure 2. Simulated datasets.** Performance on simulated mouse datasets in the presence of isotropic noise. Top: performance of different classifiers on data where each of the 295 strains is represented by an equal number of reads. Bottom: performance of the same when strain abundance is determined by counts from 16S data.

**Figure 3. Genome features.** Unique features of genomes picked up in different ways. A) Weight distributions as learned by the Autocross neural network for a general-purpose dataset. B) Comparison of the average peptide pair counts per gene for select strains, demonstrating the variety visible between them. D) Codelta comparisons. Three Firmicute strains, showing strong taxonomic correlation in their similarity. Cyan cells show a positive correlation between pairs of amino acid dimers, whereas yellow cells show a negative correlation. E) Codelta graph for random genomes with the same GC content as the *Candidatus Carsonella rudii, E. coli,* and *M. phosphovorus* genomes, illustrating the relationship between sequence complexity and environment.

**Figure 4. Real data. A)** Results of classifying the actual datasets collected from the colons of non-obese diabetic (NOD) mice comparing classification of Gist, KRAKEN, NBC, and CLARK against one another. Each community consists of 9 strains (altered Schaedler flora, or ASF). Data analysis was approximated with the same strains from Figure 2, which included only one of the ASF strains. B) Genera found during dataset classification for Gist and NBC.

